# Robust identification of deletions in exome and genome sequence data based on clustering of Mendelian errors

**DOI:** 10.1101/209478

**Authors:** Kathryn B. Manheimer, Nihir Patel, Felix Richter, Joshua Gorham, Angela C. Tai, Jason Homsy, Marko T. Boskovski, Michael Parfenov, Elizabeth Goldmuntz, Wendy K. Chung, Martina Brueckner, Martin Tristani-Firouzi, Deepak Srivastava, Jonathan G. Seidman, Christine E. Seidman, Bruce D. Gelb, Andrew J. Sharp

## Abstract

Multiple tools have been developed to identify copy number variants (CNVs) from whole exome (WES) and whole genome sequencing (WGS) data. Current tools such as XHMM for WES and CNVnator for WGS identify CNVs based on changes in read depth. For WGS, other methods to identify CNVs include utilizing discordant read pairs and split reads and genome-wide local assembly with tools such as Lumpy and SvABA, respectively. Here, we introduce a new method to identify deletion CNVs from WES and WGS trio data based on the clustering of Mendelian errors (MEs). Using our Mendelian Error Method (MEM), we identified 127 deletions (inherited and *de novo*) in 2,601 WES trios from the Pediatric Cardiac Genomics Consortium, with a validation rate of 88% by digital droplet PCR. MEM identified additional *de novo* deletions compared to XHMM, and also identified sample switches, DNA contamination, a significant enrichment of 15q11.2 deletions compared to controls and eight cases of uniparental disomy. We applied MEM to WGS data from the Genome In A Bottle Ashkenazi trio and identified deletions with 97% specificity. MEM provides a robust, computationally inexpensive method for identifying deletions, and an orthogonal approach for verifying deletions called by other tools.

## Introduction

Structural variation (SV), particularly *de novo* deletions, has been implicated in many human diseases including autism spectrum disorders, developmental delay, schizophrenia and congenital heart disease (Weischenfeldt et al., 2013; Gilissen et al., 2014; Glessner et al., 2014; Szatkiewicz et al., 2014; Brandler et al., 2015). Previously identified using microarrays, many tools have been developed in the past ten years to identify SV from next generation sequencing (NGS) data (Tattini et al., 2015). These tools utilize three main lines of evidence to detect SV: changes in read depth, discordant read pairs and split reads. Assembly methods including genome-wide local assembly and *de novo* assembly are also available (Weisenfeld et al., 2014; Wala et al., 2017).

With respect to whole exome sequencing (WES) data, one tool to identify copy number variants (CNVs) is XHMM, which identifies changes in normalized read depth within a cohort (Fromer and Purcell, 2014). Although widely used for identifying CNVs from WES data, XHMM has several limitations, including a minimum cohort size and the requirement that CNVs must include at least three exons. Typically, ~20% of putative CNVs identified by XHMM fail to be confirmed, and its sensitivity is limited (Glessner et al., 2014). For example, one study that used both XHMM and SNP arrays to identify *de novo* CNVs found that XHMM failed to detect 63% of CNVs identified by the SNP array(Glessner et al., 2014). The limited sensitivity of XHMM stems from the limitations of WES, some of which can be overcome with whole genome sequencing (WGS).

Multiple tools have been developed to identify SV from WGS data including CNVnator and Lumpy (Abyzov et al., 2011; Layer et al., 2014). While CNVnator identifies CNVs based on changes in normalized read depth (Abyzov et al., 2011), Lumpy utilizes discordant read pairs and split reads to identify deletions, duplications and other types of SVs (Layer et al., 2014). Lumpy is often used in combination with CNVnator to take into account changes in read depth. In order to estimate the sensitivity and false discovery rate (FDR), SVs identified by CNVnator and Lumpy were both compared to SVs identified in the 1000 Genomes Project by other SV callers (*e.g*., Delly, Pindel). Although both tools are reported to have a low FDR (0.4 – 3%) and high sensitivity (60 – 90%) (Abyzov et al., 2011; Layer et al., 2014), the accuracy of these tools diminishes when used for identifying *de novo* SV (Kloosterman et al., 2015). This problem results from a lack of sensitivity when identifying SVs: false negatives in parental samples lead to a high false positive rate for calling *de novo* SV, creating a significant challenge when attempting to identify *de novo* events that are potentially pathogenic.

Here, we describe a novel approach called the Mendelian Error Method (MEM) to identify and/or validate deletion SV in trios with WES and WGS data. MEM is based on the principle described in McCarroll *et al*. 2006 (McCarroll et al., 2006), where the presence of a heterozygous deletion reduces the underlying genotype to a hemizyous state. As genotype callers such as GATK assign diploid genotypes to autosomal loci,regions of heterozygous deletion are erroneously assigned homozygous genotypes. In the context of a trio design, variants within heterozygous deletions frequently display Mendelian errors as a result of this genotype mis-assignment (illustrated in Figure 1). We, therefore, hypothesized that clusters of Mendelian errors could be used as a robust signal for the presence of underlying deletions in sequencing data from trios. We applied MEM to both WES and WGS trio data from the Pediatric Cardiac Genomic Consortium (PCGC) and compared results to deletions identified by XHMM, CNVnator and Lumpy. Overall, our results show that MEM identifies both inherited and *de novo* deletions with a positive predictive value (PPV) exceeding 90%, and identifies additional *de novo* deletions that are missed by other SV callers.

**Figure 1:**
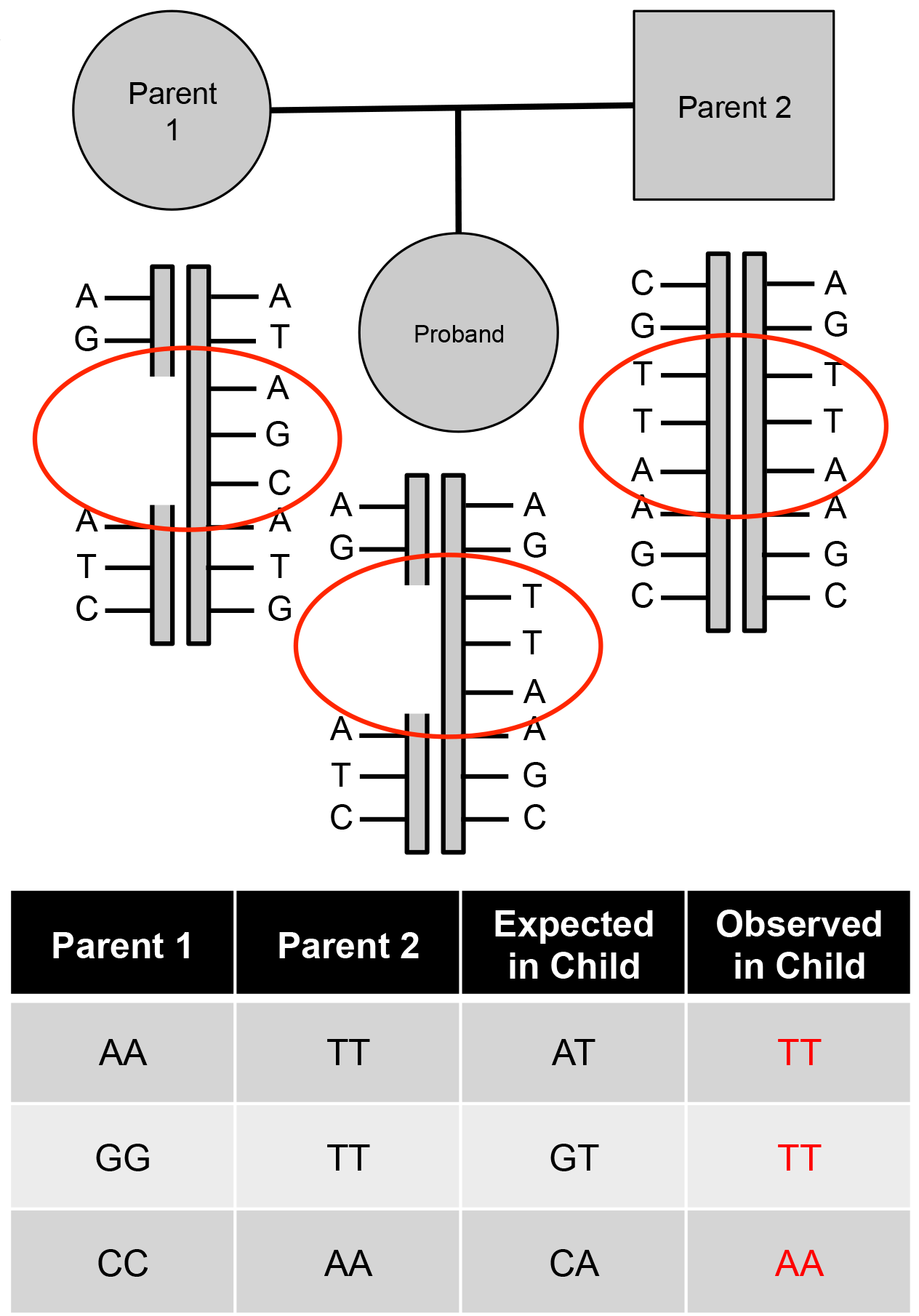
Schematic of MEM principle. Diagram of trio where proband inherited a deletion from parent 1. Tools report homozygous genotypes (red) that violate Mendelian laws of segregation in the case of hemizygosity due to a heterozygous deletion. Adapted from McCarroll *et al.* 2006 (McCarroll et al., 2006).

## Methods

### WES and WGS in cases with CHD

Probands were recruited from 10 centers in the United States and United Kingdom as part of the Congenital Heart Disease Genetic Network study of the PCGC as described previously (Homsy et al., 2015). Cases (n=2,601) were subject to WES at the Yale Center for Genome Analysis as described previously (Homsy et al., 2015), with a mean depth of 107x. All genomic coordinates quoted are based on human genome hg19/build 37. Variants were called following the n+1 protocol from GATK.

Three hundred and fifty probands and their parents from the PCGC were selected for WGS; of note 332 also have WES data. Cases were sequenced at the Broad Institute(n=25), New York Genome Center (n=25) and Baylor College of Medicine Human Genome Sequencing Center (n=300). Samples were sequenced with PCR-free library preparation (n=325) or with SK2-IES (n=25) to a mean depth of 30x on Illumina HiSeq X Ten sequencers. Variants were called by GATK HaplotypeCaller (version 3.3.2) following GATK best practices for n+1 joint calling (https://software.broadinstitute.org/gatk/best-practices/).

### WES and WGS of healthy population cohort

Trios representing a typical population cohort (n=1,683) were provided by the Simons Foundation Autism Research Initiative Simplex Collection. Simplex families (two unaffected parents, one child with autism spectrum disorder, and one unaffected sibling) underwent WES using DNA extracted from peripheral blood cells, with a mean depth of 117x (O’Roak et al., 2011; Sanders et al., 2012; Iossifov et al., 2014). Trios of unaffected siblings and parents served as a typical population cohort for comparison.

Five hundred and nineteen quartet families selected from the Simons Simplex Collection (SSC) underwent WGS at the New York Genome Center. Samples were sequenced with either a PCR-based library preparation on an Illumina Hi-Seq 2000 (n=39) or PCR-free library preparation on an Illumina HiSeq X Ten (n=480). Sequencing was performed with 150-bp paired reads with median coverage of 37.8x per individual. Detailed information regarding this cohort can be found in Werling *et al.* (Werling et al., 2017)

Variants were called using GATK HaplotypeCaller (version 3.1-1-g07a4bf8, n=19, version 3.2-2-gec30ce, n=21, version 3.4-0-g7e26428, n=479). GATK best practices (https://software.broadinstitute.org/gatk/best-practices/) were followed. Trios comprising an unaffected sibling and their parents were used as a typical population cohort for comparison in this study with permission from the SSC.

### Genome in a Bottle (GIAB) WGS with Illumina

The GIAB Ashkenazi Jewish (AJ) trio was subject to WGS using both short and long read methodologies. 148-bp paired-end reads were generated with an Illumina Hiseq instrument. Reads were aligned with BWA-mem (details in Zook *et al*., 2016) (Zook et al., 2016). Variants were called by GATK HaplotypeCaller (version 3.3.2) following GATK best practices using n+1 joint calling.

### GIAB deletions for AJ trio

GIAB provided draft benchmark structural variants (SVs) for the AJ trio (v0.3.0a). SVs from 119 different tools were compared and merged using the tool SURVIVOR (Jeffares et al., 2017), which required the breakpoints to be within 1000 bp. Deletions identified by a minimum of two tools were compared to deletions identified by MEM using bedtools and required a 20% reciprocal overlap.

### Mendelian Error Method (MEM) Pipeline – Figure 2

#### 1. Extract Mendelian errors (MEs) from WES and WGS VCFs

MEs were extracted based on genotypes reported in the joint VCF produced by GATK best practices, using in-house perl scripts or vcftools. Table S1 includes the eight scenarios considered as MEs that could represent a deletion.

**Figure 2:**
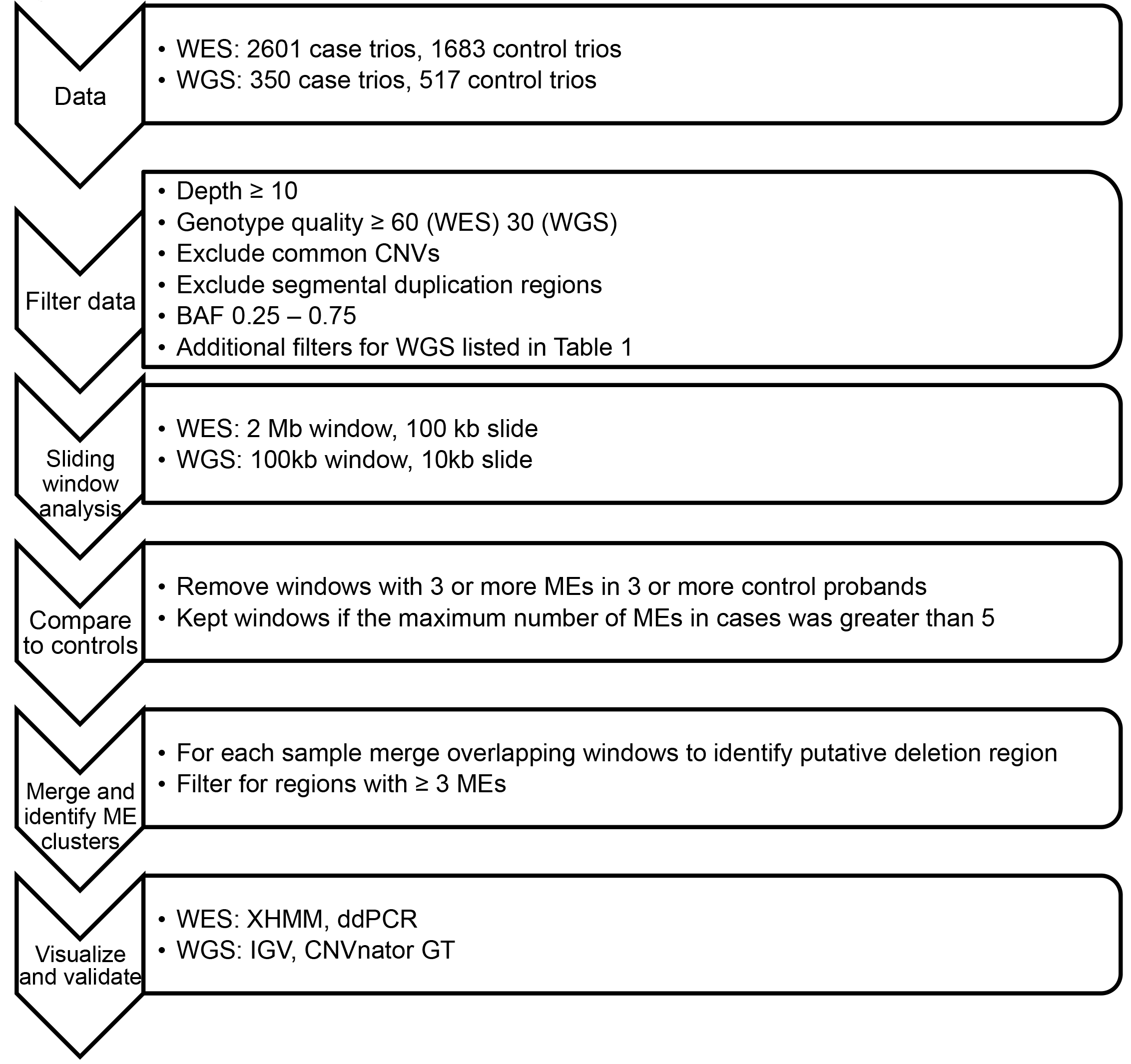
MEM pipeline for WES and WGS data.

#### 2. Filtering

Variants in PCGC, GIAB and SSC trios were filtered using the following criteria: read depth ≥ 10, genotype quality >60 for WES and >30 for WGS (Table S2). B allele frequency (BAF, defined as the alternate allele depth/total depth) was calculated for heterozygous SNVs, and those with a BAF <0.25 or >0.75 were excluded. Regions overlapping segmental duplications obtained from the UCSC Genome Browser track were excluded. CNVs with a minor allele frequency ≥0.05 in European, African or East Asian ancestry as identified in Conrad *et al*. were excluded (Conrad et al., 2012). For WGS, SNVs with a mappability score <1 were excluded, based on the UCSC Genome Browser track “Alignability of 100mers by GEM from ENCODE/CRG(Guigo)”. Regions with tandem repeats, taken from the UCSC Genome Browser track “Simple Repeats” and expanded ±5 bp, were excluded. The Hardy Weinberg equilibrium (HWE) statistic was calculated using vcftools for SNVs with a minimum allele frequency of 0.01 in parents. Any SNVs with a HWE p-value equal to zero were removed.

#### 3. Sliding window analysis

We generated 2-Mb windows with 95% overlap for WES analysis and 100-kb windows with 90% overlap for WGS analysis using Bedtools (version 2.26.0) makewindows. In house bash scripts utilizing Bedtools intersect were used to calculate the number of MEs for each window. This was applied to each sample in the PCGC and SSC cohorts separately.

For each unique window, the number of probands with MEs, the minimum number of MEs, the maximum number of MEs and the average number of MEs per proband were calculated for PCGC and SSC probands. We filtered for windows where the average number of MEs per proband was >2 MEs.

#### 4. Comparison to population cohort

Windows with MEs in PCGC cases were compared to corresponding windows in the SSC population cohort. Windows with a ME cluster in three or more SSC probands were excluded, except if the maximum number of MEs in cases was >5.

#### 5. Merge windows

For each sample overlapping windows with MEs were merged to identify putative deletion regions. The minimum, maximum and average number of MEs per window was calculated for each region. The number of MEs in each putative deletion region was calculated in SSC probands and regions with ME clusters as described in Step 4 were removed from further analysis.

#### 6. Filter for ME clusters

Finally, we filtered for regions with an average number of MEs per window >2 in cases. We identified the first and last ME within each region and used these as the coordinates for the putative deletions.

### Visualization

#### 1. XHMM

For putative deletions identified with MEM from the PCGC WES cohort, we extracted zF scores of the PCA-normalized read depth for each exon from XHMM (Fromer and Purcell, 2014). Putative deletions were inspected visually (Figure S1) and exons with zP scores <-2 were considered candidates for deletions.

#### 2. IGV

Integrated Genomics Viewer (IGV, version 2.3.34) pileup visualization was used as one method for deletion validation. Variants were visualized in the proband and parents. Deletions were excluded if any of the following aspects were detected: multiple reads with quality scores of zero in child or parents, no clear drop of coverage in the proband, or the presence of heterozygous SNVs in the proband.

### CNVnator

CNVnator identifies CNVs in WGS data based on changes in normalized read depth (Abyzov et al., 2011). Deletions were called for each case proband and the GIAB proband with CNVnator (version 0.3.2) and genotyped for putative copy number within the CNV regions on a scale from 0 – 3. We considered scores between 0.7 – 1.4 as indicating a heterozygous deletion. *De novo* deletions were identified by filtering for a score <1.4 in the child and >1.4 in the parents. We overlapped putative deletions in WGS cases identified using MEM with *de novo* deletions identified by CNVnator using Bedtools intersect, requiring a 25% reciprocal overlap. In the AJ trio, we overlapped putative deletions identified with MEM with both inherited (proband genotype <1.4) and *de novo* deletions called by CNVnator, and considered all intersections with at least 1 bp of overlap.

### Lumpy

Lumpy identifies SVs based on discordant read pairs and split-reads (Layer et al., 2014). Deletions were called for each case proband and the GIAB proband with Lumpy (version 0.2.13) and genotyped using SVtyper (version 0.0.4). *De novo* deletions were identified based on proband and parent genotypes. We overlapped PCGC WGS MEM deletions with Lumpy *de novo* deletions in the same manner as CNVnator. In the AJ trio, we overlapped putative deletions identified with MEM with both inherited and *de novo* deletions by Lumpy, and considered all intersections with at least 1 bp of overlap.

### SvABA

Deletions were called with SvABA from 350 WGS trios based on genome-wide local assembly (Wala et al., 2017). Default parameters were employed to identify putative copy number variants, which were further validated by IGV visualization prior to digital droplet PCR analyses.

### Deletion validation

Digital droplet PCR (ddPCR) was used to validate MEM WES deletions and WGS *de novo* deletions identified by CNVnator and Lumpy, as previously reported (Mazaika and Homsy, 2014) with the following modification. PCR primers that amplified a portion of the putative CNV were designed to avoid homopolymer runs or probes that begin with G. PCR-positive droplets were identified by EvaGreen dye (DNA-bound emission at 500/533 nm). CNV product positive droplets were EvaGreen dye positive, VIC negative. A VIC probe targeting the RPP30 gene was used as reference. Reaction mixtures of 20μL volume comprising ddPCR Master Mix (Bio-Rad), relevant forward and reverse primers and probe(s) and 50ng of DNA were prepared, ensuring that <40% of the 5000-10000 droplets ultimately produced were positive for Evagreen dye and/or VIC signal. For *de novo* CNV confirmations, DNA from the subject with CHD and parents was used. After thermal cycling, plates were transferred to a droplet reader (Bio-Rad) that flows droplets single-file past a 2-color fluorescence detector. Differentiation between droplets that contain target and those that did not was achieved by applying a global fluorescence amplitude threshold in QuantaSoft (Bio-Rad). The threshold was set manually based on visual inspection at approximately the mid-point between the average fluorescence amplitude of positives and negative droplet clusters on each of the EvaGreen dye and VIC channels. Confirmed CNV duplications had ≈50% increase in the ratio of positive to negative droplets, as did the reference channel. Conversely, confirmed CNV deletions had approximately half the ratio of positive to negative droplets, as did the reference channel. CNVs that were called, but were unable to be confirmed or rejected due to ddPCR technical failure or DNA unavailability were excluded from analysis.

## Results

### MEM identifies inherited and de novo deletions from WES trios

The MEM pipeline was used to analyze WES data from 2,601 PCGC trios and 1,683 healthy trios from the SSC. Windows with ME clusters in SSC probands were removed as described in Methods in order to limit our findings to those of likely relevance to the pathogenesis of congenital heart disease (CHD). MEM identified a final set of 171 merged and filtered regions containing putative deletions in the PCGC probands (Table S3). We used the location of the first and the last ME in each region with a ME cluster to define the minimal coordinates for the deletion. We utilized XHMM read depth data to perform an initial assessment of the accuracy of our MEM deletion calls. The proband’s normalized XHMM z-scores for each exon within the deletion identified by MEM were compared to the rest of the cohort (Figure S1). The presence of outlier negative zc scores in the proband suggested a deletion. The parents’ z-scores were also compared to the rest of the cohort to determine if the deletion was inherited or *de novo.* In this manner, 58 deletions were determined to be *de novo*, and 79 were noted to be inherited. Of note, the exons in 13 ME clusters did not have negative normalized zi scores, and seven ME clusters showed inconsistent scores, with some exons showing reduced XHMM z-scores, while other exons were within the normal range (z-score >-2), suggesting that these 20 calls could be false positives.

We directly compared the performance of MEM for the detection of *de novo* deletions with that of XHMM. Fifty deletions were called by both tools, 46 by XHMM alone, and 25 by MEM alone (Figure 3A). Of note, the 25 MEM-exclusive deletions included 13 that showed no reduction in z-scores with XHMM for proband or parents and, thus, could represent either *de novo* deletions or false positives. We considered the size of the deletions that MEM did and did not identify. For deletions ≥200 kb, MEM identified 100% of deletions, however for deletions <200 kb MEM identified 24% of deletions (Figure 3B). The 46 XHMM-exclusive deletions had a mean size of 35 kb and, therefore due to an insufficient number of SNPs within them, could not be identified by MEM with high recall.

**Figure 3:**
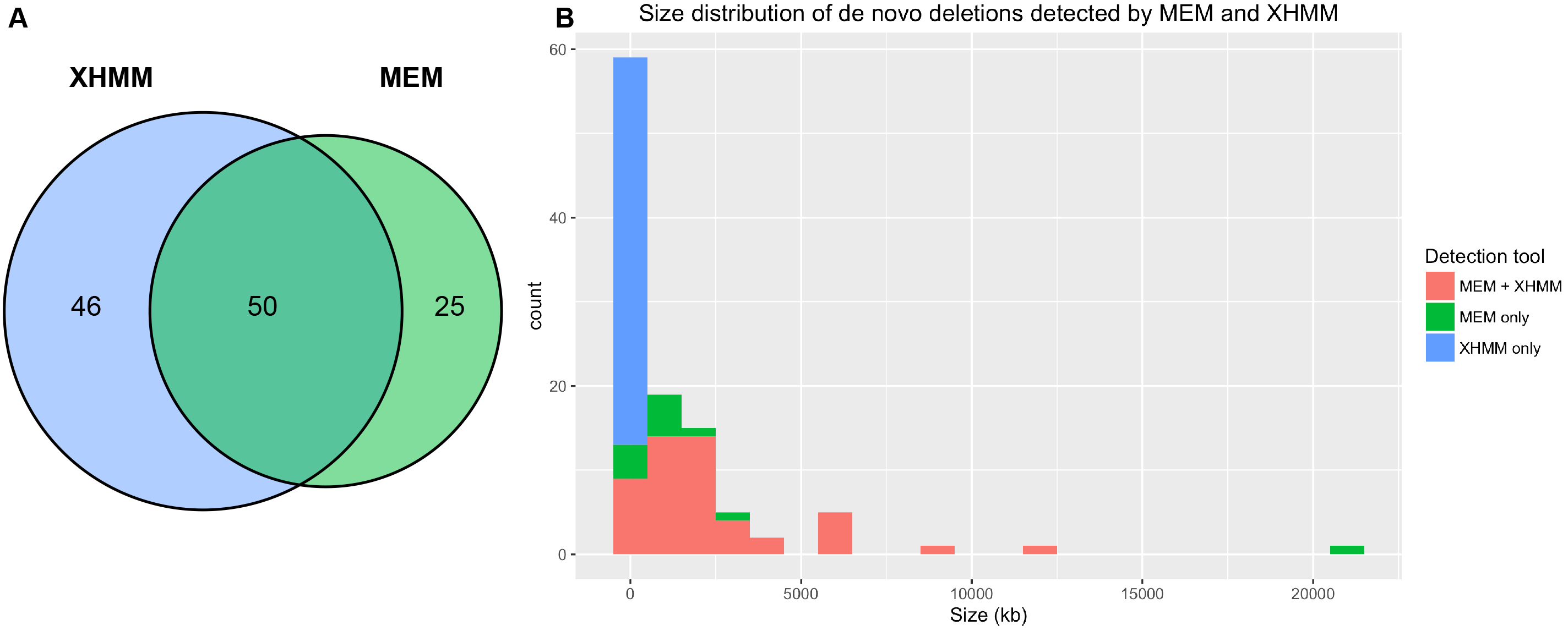
A) Comparison of *de novo* deletions called by XHMM and MEM. B) Size distribution of *de novo* deletions called by XHMM. Colors in stacked histogram indicate which tools detected the deletion (red = MEM and XHMM detected, green = MEM detected and not XHMM, blue = XHMM detected and not MEM).

From the 171 MEM deletions, 36 overlapped with deletions previously confirmed by digital droplet PCR (ddPCR). For the remaining 135 deletions, we performed ddPCR, which was successful for 109 deletions. Ninety-six out of 109 were confirmed as true deletions, achieving a positive predictive value (PPV) of 88.1%. Surprisingly, the results from ddPCR indicated that five of the regions with the ME cluster were inherited duplications. Thus, overall 137/145 (94.5%) of ME clusters identified by MEM were confirmed as true CNVs. Deletions identified as inherited by inspection of XHMM zc score plots confirmed with a PPV of 86% (49/57 inherited, 3/57 *de novo*). From the possible false positives, two out of eight deletion regions without negative normalized zp scores in XHMM were confirmed, and four of six regions with inconsistent loss of exons confirmed. Finally, 26 *de novo* deletions were confirmed, four exclusively identified by MEM.

### Enrichment of deletions on chromosome 15q11.2

With MEM, we identified 15 deletions (13 inherited, 2 *de novo*) ranging from 11 kb to 1 MB in the chromosome region 15q11.2 in PCGC probands. These deletions fall in a known microdeletion region between breakpoints (BP) 1 and 2, with a population frequency of 0.25% (Cafferkey et al., 2014). Deletions in this region occurred at a frequency of 0.58% (15/2,601) in the PCGC cohort, and are therefore enriched compared to the reported population frequency (binomial, p=0.004) and to SSC probands, which had a deletion frequency of 0.24% (4/1,683) deletions in this region (binomial, p=0.002).

### Identification of uniparental disomy (UPD) in WES trios by MEM

Following ME extraction and applying quality filters (Table S1), the majority of trios had between 0.6 – 2% of loci that were scored as MEs (Figure 4A). We identified eight probands with an elevated rate of MEs distributed across an entire chromosome, suggestive of possible uniparental disomy (UPD). Prior microarray experiments noted UPD of chromosome 15 for one proband, and an extended region of homozygosity on chromosome 16 for a second proband. However, there was no prior indication of UPD in the other six cases.

**Figure 4:**
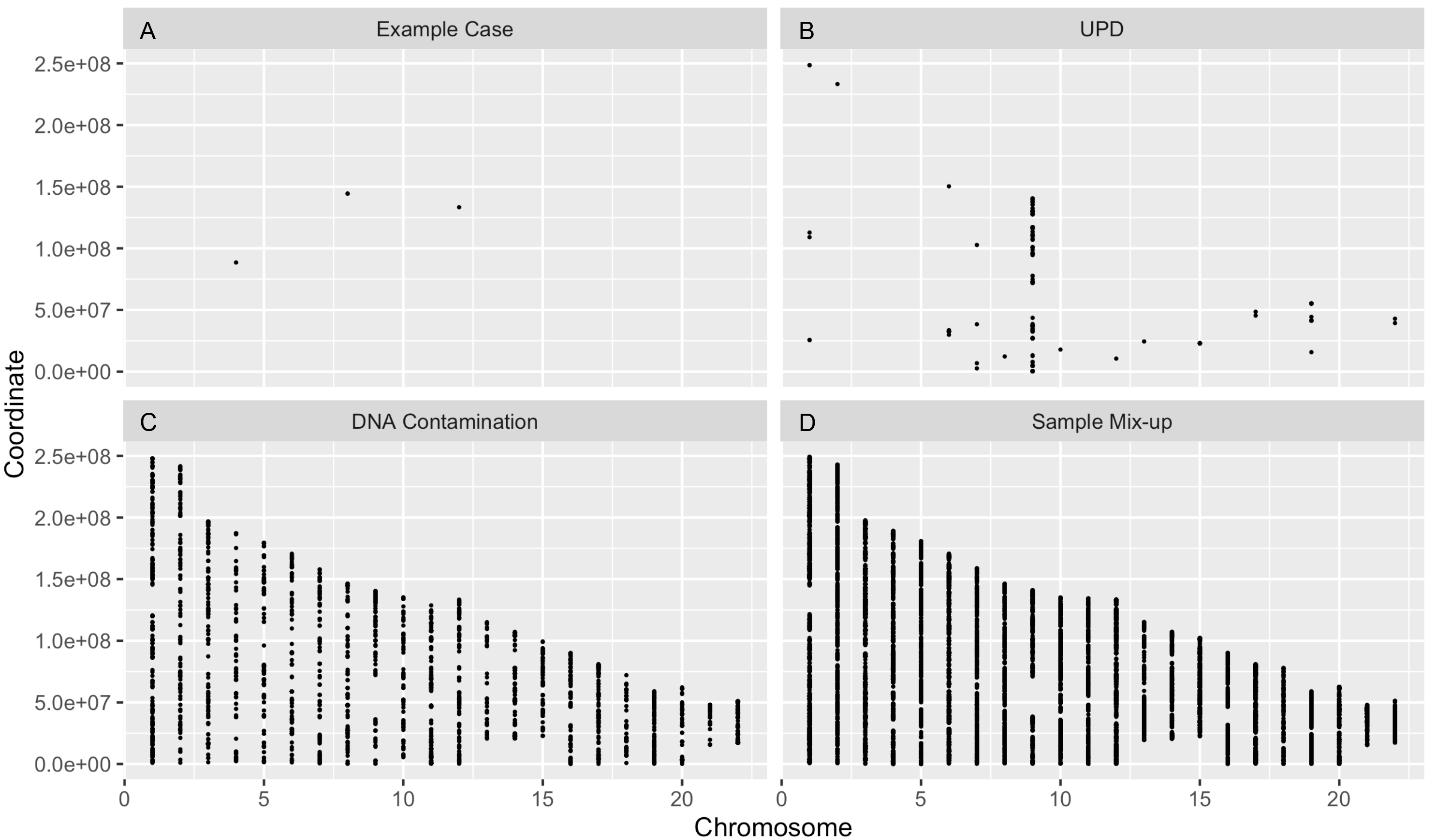
MEs plotted by chromosome. A) MEs in a trio after quality filtering. B) Sample with UPD on chromosome 9. C) Trio with DNA contamination. D) Trio with a sample mix.

All eight instances of UPD were classified as maternal heterodisomy, based on the presence of heterozygous maternal SNPs. The heterodisomic inheritance was for chromosomes 4 (x2), 8, 9, 14, 15 and 16 (x2). UPD was not found in any SSC probands, and was therefore enriched in cases (binomial, p=0.026).

### MEs identify irregularities in WES trios

We identified two other distinct ME patterns that were informative. Twenty trios had a dramatically higher rate of MEs (~50% of all SNVs), which were distributed across every chromosome (Figure 4D). Nearly all of the MEs were attributable to lack of inheritance from one parent, suggesting either a sample switch or incorrect paternity.

Similarly, we observed an elevated, but lower, rate (20-30%) of MEs distributed across the entire genome in six other probands (Figure 4C). We hypothesized that this pattern might be due to DNA contamination, which was confirmed with the program VerifyBamID (Jun et al., 2012).

All samples with likely sample mix-ups, DNA contamination or UPD were excluded from further analysis.

### ME clusters are non-random in the genome

Before applying MEM to WGS data, we first needed to determine if the increased SNV density in WGS data relative to WES data could lead to ME clusters by chance alone. To test this, we generated a null model of SNV clusters across the genome. We only considered heterozygous SNVs, and also applied additional filters for genotypes generated from WGS as shown in Table 1. After applying these quality filters, the median number of MEs per proband among the 350 PCGC WGS trios was 317. We then ran 1000 permutations of selecting 317 informative SNV positions from one trio, assuming those were MEs, and implemented MEM with a 100-kb window and 10-kb slide. We calculated the number of windows with SNV clusters divided by the number of windows with at least 1 SNV. The null model had a mean of 0.3% of windows with a SNV cluster (Figure S2). In contrast, 21.4% of windows with at least 1 ME among the PCGC WGS probands had a ME cluster and they were infrequent across the genome (Figure S2). From these results, we inferred that ME clusters in WGS were likely non(random and were likely identifying underlying deletions.

**Table 1:**
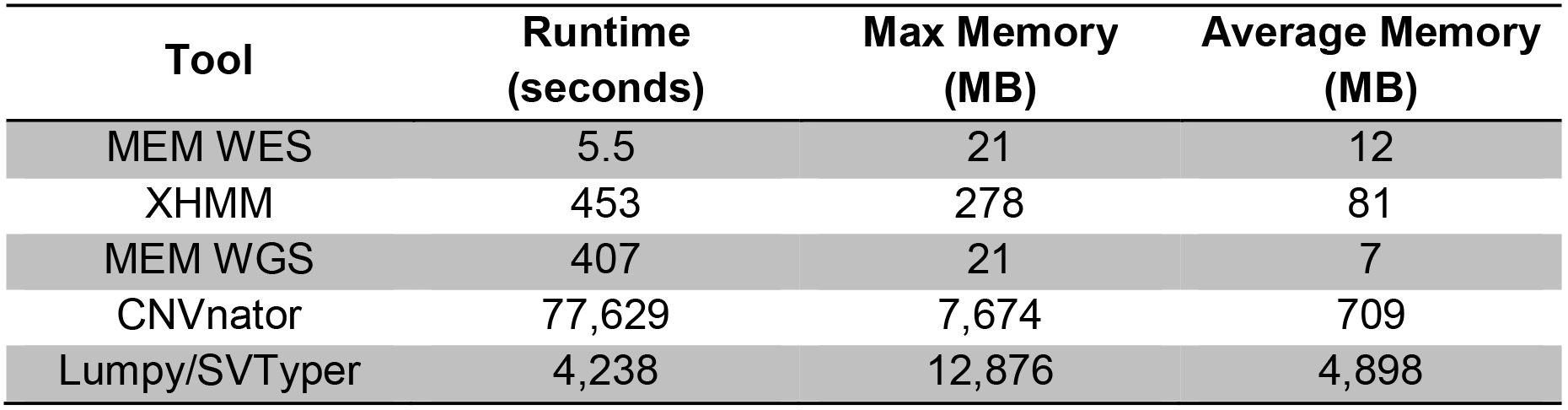
Computational resources required for NGS CNV detection tools

### Mendelian error clusters identify deletions from GIAB Ashkenazi trio

To test the robustness of MEM for calling deletions from WGS, we identified putative deletions using MEM based on genotypes generated using Illumina short read WGS data for an Ashkenazi Jewish (AJ) trio sequenced by the GIAB consortium (Zook et al., 2016). We processed filtered SNV genotypes from the Illumina WGS data in this trio using the parameters listed in Table 1 and searched for ME clusters. Using the MEM pipeline we identified 32 putative deletions (Table S3) that contained an average of 9.4 MEs, with a mean size of 31.5 kb.

To determine the accuracy of the MEM deletion calls, we intersected them with draft benchmark deletions provided by GIAB. Requiring a 20% reciprocal overlap between deletions, 27/32 MEM deletions overlapped with those from GIAB. After removing the 20% overlap requirement 31/32 MEM deletions overlapped. The five deletions that did not overlap by 20% were visualized in IGV, where we found evidence for a deletion in 4/5. Therefore, MEM identified deletions with 97% precision from WGS for the GIAB AJ proband. Of note, one 215-kb MEM deletion overlapped two GIAB deletions. Visualization in IGV confirmed the presence of two separate deletion events at this locus, which the distribution of MEs also supports (Figure S3).

Next, we looked at the deletions identified by GIAB that MEM did not identify (n=24,090). These do not include deletions in segmental duplication regions but do include 14,690 deletions at tandem repeat loci. Due to the challenges of sequencing tandem repeats with short read sequencing we would not expect MEM to accurately identify deletions with tandem repeats, as variant calling is unreliable in these regions. The MEM false negatives (FNs) had a median size of 39 bp and a mean size of 306 bp and were attributable to inadequate number of MEs in those deletions as 93.5% did not include any MEs before filtering. Only 1% of the MEM FNs were related, at least in part, to the filtering of MEs, having >2 MEs prior to filtering.

We also compared the MEM calls for the AJ trio to calls from CNVnator and Lumpy. Of the 32 MEM deletion calls, 27 (84%) and 23 (72%) overlapped with calls from CNVnator and Lumpy, respectively. There were many calls from CNVnator and Lumpy that were not made by MEM, however most of them contained no MEs. ME filtering accounted for 21% of FNs from CNVnator calls and 6% of FNs from Lumpy calls.

### MEM identifies deletions from WGS trios

Based on the promising results from GIAB, we proceeded to apply the MEM pipeline to identify deletions from 350 WGS case trios from the PCGC, and 517 healthy trios from the SSC. From the PCGC trios, MEM identified 6,645 regions with ME clusters (mean=19.1/proband) that ranged in size from 3 bp to 9 Mb, with a median size of 2.9 kb and a mean size of 20 kb (Table S3). Eleven percent of regions included exons. We used the first and last MEs as coordinates for the putative deletions. For 332 PCGC trios that have both WES and WGS data we compared the deletions identified by MEM from both data sets. MEM identified 11 deletions from WES, all of which were detected by MEM with WGS. All of the deletions were the same size or larger when detected by WGS except for one. This is expected as the increased SNP density of WGS provides more informative sites for MEM, thus facilitating a better estimate of the deletion size.

To determine if the ME clusters in WGS data identified true deletions, we integrated normalized read depth data from CNVnator. Each region was labeled with a CNVnator score where 0 corresponds to a homozygous deletion, 0.7-1.5 to a heterozygous deletion, 1.5-2.4 to being normally diploid and >2.4 to a duplication. The vast majority (97%) of MEM deletions had a CNVnator score between 0.7 – 1.5 suggesting MEM was identifying true heterozygous deletions (Figure S4). We visualized MEM deletion calls with a CNVnator score >1.5 in IGV. Based on this manual curation, we concluded that the majority (66%) were false positives, but 34% were heterozygous deletions: 10% covering the entire region and 24% being either a deletion of a portion of the region or two smaller deletions located close together. In addition, we visualized in IGV a test set of MEM deletions with a range of CNVnator scores. The vast majority of false positives (93.5%) had a score of 1.5 or greater, while 100% of the true or possible deletions had a score between 0.7 and 1.5 (Figure S5). Overall, our comparison with read depth data supports a PPV of 92% (Supplementary Formula 1) for identifying heterozygous deletions from WGS with MEM.

Next, we identified which MEM deletions were *de novo* based on the proband and parents’ CNVnator scores. We used two sets of filters (Table S4) and identified 37 putative *de novo* deletion calls (mean = 0.12 *de novo* deletions/proband) After visualization in IGV, we determined that 20/37 represented likely true *de novo* deletions, while 17 were inherited. We compared these to *de novo* deletions identified by CNVnator, Lumpy and a third WGS tool called SvABA that uses genome-wide local assembly to identify SV (Wala et al., 2017). The deletions called by the other SV tools were confirmed by ddPCR. Of the 20 *de novo* deletions found by MEM, five were also identified by CNVnator, Lumpy, and SvABA, three were identified by CNVnator and SvABA but not Lumpy, and 12 were not found by the three other tools. Thirteen additional *de novo* deletions were identified with a combination of CNVnator, Lumpy and SvABA: all three tools but not MEM (n=7), CNVnator and SvABA (n=2), CNVnator and Lumpy (n=1), CNVnator only (n=2), and SvABA only (n=1). None of these deletions, which had a median size of 6.5 kb, included any MEs, suggesting MEM is less sensitive for deletions smaller than ~10 kb in WGS.

### MEM is computationally efficient

We compared the computational resources required for MEM and the other CNV detection tools used in this study for deletion identification in one trio (Table 1). Runtime and memory for all tools were based on the use of an Intel Haswell 2.4 GHz processor with 64 GB memory and Cray nodes. We did not utilize parallelization for any of the tools. Runtime and memory for MEM was calculated for Step 1 of the MEM pipeline (ME extraction). All other steps in the MEM pipeline can be performed on the command line and do not require significant time or memory. Of note, resources required for the preliminary steps for all tools (DepthOfCoverage for XHMM, Samblaster for Lumpy, and variant calling for MEM) were not included.

For WES, MEM required 5.5 sec and an average of 12 MB of memory per trio. XHMM required 453 sec and on average 81 MB of memory. For WGS, MEM required 407 sec and an average of 7 MB of memory per trio. CNVnator required 77,629 sec and, on average, 709 MB of memory. Lumpy/SVTyper required 4,238 sec and an average of 4,898 MB of memory. SVTyper produced genotypes for deletions only and not other types of SV (duplications, translocations, inversions). For both WES and WGS, MEM performed significantly faster and required significantly less memory compared to other CNV detection tools. Of note, ME extraction execution time grows sub-linearly based on the number of trios present in the VCF, however average memory required does not increase significantly.

## Discussion

A variety of tools have been developed to identify CNVs including XHMM and CoNIFER for WES, and CNVnator, Lumpy and SvABA for WGS. Each of these tools has limitations such as a requirement for 50 samples, the need for extensive computational resources, or that up to 20% of CNVs will fail to confirm. In addition, false negative calls in parents lead to a high false positive rate for *de novo* deletion CNV calls, making the identification of true *de novo* CNVs difficult and time intensive. As documented in this report, we developed a novel method, MEM: the Mendelian Error Method, to identify deletion CNVs based on ME clustering. This orthogonal method identifies deletions with a PPV >90% for both WES and WGS, and identifies additional *de novo* deletions compared to other SV callers.

When used with WES, we demonstrate that MEM has several advantages compared to XHMM. First, MEM can be used on a single trio, while XHMM requires a minimum of 50 samples to accurately normalize read depth and calculate z-scores. Second, MEM requires substantially less memory and runtime compared to XHMM. Third, MEM can be used as a method for quality control, as it can identify UPD, sample mix-ups and DNA contamination. MEM is also a worthwhile complementary tool to XHMM as MEM identified additional *de novo* deletions that XHMM missed due to spurious evidence of inheritance or seemingly inconsistent loss of exons. In addition, MEM identified deletions with less than 3 exons with high precision, albeit with low sensitivity. The combination of evidence from both XHMM and MEM can increase our ability to identify smaller deletions with high precision and increased sensitivity, as well as reducing the need for PCR-based validation, which is expensive and time-consuming.

CNV identification from WGS data is still under development. We propose MEM as a worthwhile addition to the WGS CNV identification toolbox as it can be efficiently implemented in less than a day and identifies deletions with a >90% PPV. It can be implemented on a large cohort without significantly increasing the computational requirements, and identifies additional *de novo* deletions compared to CNVnator, Lumpy and SvABA. While there are other SV tools for WGS data (*e.g*., Delly, Pindel), the methods utilized by CNVnator, Lumpy and SvABA, represent three primary ways to identify CNVs: changes in read depth, discordant/split reads and local assembly, yet MEM identified additional *de novo* deletions. Equally helpful is the orthogonal nature of MEM, which may reduce the need for PCR validation for deletions identified by MEM and a second tool.

MEM’s primary limitation is the need for a complete trio, as many cohorts only recruit singletons. The trio design is necessary in order to identify MEs and, therefore, cannot be avoided. MEM is also limited regarding the size of the deletions it can detect with high recall, which is a function of the SNV density in NGS data. WES deletions <200 kb are identified with 24% recall, while deletions >200 kb are identified with 100% recall. Of note, although the smaller deletions are not identified with high sensitivity, the PPV remains high when they are called (78%). Based on deletions identified in GIAB AJ trio, MEM identifies deletions from WGS with a range of sizes (100 – 660,000 bp); however, we estimate that MEM has ~1% recall for deletions smaller than 3 kb and only 18% recall for deletions 3-10 kb. Deletions >10 kb are identified with 45% recall. For this reason MEM applied to WGS is particularly valuable as a secondary and orthogonal method to confirm deletions identified by other tools, as the PPV is 92-97% with WGS data.

MEM’s sensitivity was also reduced by ME filtering, which accounted for ~5% of the false negatives. Filtering is necessary in order to remove MEs caused by poor genotyping or other errors and to achieve a high PPV. We suggest noting the number of filtered MEs when verifying deletions identified with other tools, as even the presence of 1 or 2 MEs after filtering is evidence for a deletion in 88% of calls (data not shown).

Interestingly, 3.5% of regions with ME clusters identified with MEM were scored as inherited duplications by ddPCR. Although ME genotypes are not indicative of a duplication, it is has been noted that some CNVs are complex events with multiple breakpoints comprising both deletions and duplications in close proximity (Quinlan et al., 2010). We hypothesize that this phenomenon likely underlies our observations, and that in these few cases the primer placement for ddPCR targeted a region of duplication rather than the deletion found my MEM.

The pursuit of disease-causing CNVs in family trios often focuses on the identification of *de novo* or rare CNVs. MEM identifies both inherited and *de novo* deletions, however one is unable to distinguish between inherited and *de novo* deletions without the use of a secondary tool that identifies deletions in parents. In order to identify rare CNVs from a large cohort, one must eliminate regions with deletions in the general population. This is included in the MEM pipeline in Steps 4 and 5. If population data are not available, one could determine the number of samples with deletions in each region identified by MEM as an alternative. Deletion regions found in multiple samples are less likely to be disease-causing.

We applied MEM to trios from the PCGC to identify deletions that are causal for CHD that had not been seen with previous studies (Glessner et al., 2014). With MEM, we identified and quantified two genetic mechanisms associated with CHD; BP1-BP2 deletions in 15q11.2 and UPD. Deletions in the region 15q11.2 BP1-BP2 account for ~0.3% of CHD cases in the PCGC cohort. Although 15q11.2 deletions are associated with a wide range of phenotypic anomalies, CHD have been reported in ~9% of carriers (Cox and Butler, 2015), which explains the presence of an inherited mutation present in both a proband with CHD and their apparently unaffected parent.

Using MEM, we also identified whole-chromosome maternal heterodisomy in ~0.3% of CHD cases in the PCGC cohort. The likely genetic mechanism for maternal heterodisomic UPDs is non-disjunction and subsequent trisomy rescue. Thus, there is a possibility that probands with UPD may be mosaic for trisomy of the UPD chromosome, and this mosaic trisomy could be the underlying cause of the probands’ CHD. UPD could also lead to CHD due to changes in methylation of imprinted genes. One example from the chromosomes affected in PCGC probands is chromosome 8, which harbors the known CHD gene *CHD7* (MIM:608892) that is maternally methylated (Joshi et al.,2016). Maternal heterodisomy would lead to hypermethylation and altered expression of
*CHD7.*

In conclusion, MEM is an orthogonal tool that identifies deletion CNVs with over 90% PPV and is a valuable addition to CNV detection pipelines for both WES and WGS. As NGS data becomes more accessible, the need to identify CNVs from WES and WGS data will only increase. This is particularly true with relation to disease causing CNVs as CNVs have been implicated in a number of different human diseases including congenital heart disease, schizophrenia, developmental delay and autism spectrum disorders. MEM helps overcome some of the challenges associated with identifying pathogenic CNVs due to limited specificity of current SV tools.

## Acknowledgments

The authors are grateful to the patients and families who participated in this research and team members who supported subject recruitment and sequencing D. Awad, C. Breton, K. Celia, C. Duarte, D. Etwaru, N. Rishman, M. Daspakova, J. Kline, R. Korsin, A. Lanz, E. Marquez, D. Queen, A. Rodriguez, J. Rose, J.K. Sond, D. Warburton, A. Wilpers and R. Yee (Columbia Medical School); B. McDonough, A. Monafo, J. Stryker (Harvard Medical School); N. Cross (Yale School of Medicince); S. M. Edman, J.L. Garbarini, J.E. Tusi, S.H. Woyciechowski (Children’s Hospital of Philadelphia); J. Ellashek and N. Tran (Children’s Hospital of Los Angeles); K. Flack, L. Panesar, N. Taylor (Univeristy College London); D. Gruber and N. Stellato (Steve and Alexandra Cohen Children’s Medical Center of New York); D. Guevara, A. Julian, M. MacNeal, C.Mintz (Icahn School of Medicine at Mount Sinai); and E. Taillie (University of Rochester School of Medicine and Dentistry). We also thank the Simons Foundation for Autism Research for the contribution of SSC exome trios. This work was supported by grants from the National Heart, Lung, and Blood Institute (NHLBI) for the Pediatric Cardiac Genomics Consortium (U01-HL098188, U01-HL131003, UM1-HL098147, UM1- HL098153, UM1-HL098163, UM1-HL098123, UM1-HL098162, UM1-HL128761, UM1- HL128711). The views expressed are those of the authors and do not necessarily reflect those of the NHLBI. Research reported in this paper was supported by the Office of Research Infrastructure of the National Institutes of Health under award number S10OD018522. This work was supported in part through the computational resources and staff expertise provide by Scientific Computing at the Icahn School of Medicine at Mount Sinai.

**Conflict of Interest:** The authors have no conflicts of interest to declare.

**Ethical compliance:** All procedures performed in studies involving human participants were in accordance with the ethical standards of the following Institutional Review Boards: Boston Children’s Hospital, Brigham and Women’s Hospital, Great Ormond Street Hospital, Children’s Hospital of Los Angeles, Children’s Hospital of Philadelphia, Columbia University Medical Center, Icahn School of Medicine at Mount Sinai, Rochester School of Medicine and Dentistry, Steven and Alexandra Cohen Children’s Medical Center of New York, and Yale School of Medicine. Informed consent was obtained from all individual participants or their parent/guardian included in this study.

## References

Abyzov A, Urban AE, Snyder M, Gerstein M. 2011. CNVnator: An approach to discover, genotype, and characterize typical and atypical CNVs from family and population genome sequencing. Genome Res 21:974–984.

Brandler WM, Antaki D, Gujral M, Noor A, Rosanio G, Chapman TR, Barrera DJ, Lin GN, Malhotra D, Watts AC, Wong LC, Estabillo JA, et al. 2015. Frequency and complexity of de novo structural mutation in autism. bioRxiv 1–19.

Cafferkey M, Ahn JW, Flinter F, Ogilvie C. 2014. Phenotypic Features in Patients With 15q11.2(BP1-BP2) Deletion: Further Delineation of an Emerging Syndrome. Am J Med Genet Part A 2:1916–1922.

Conrad DF, Pinto D, Redon R, Feuk L, Gokcumen O, Zhang Y, Aerts J, Andrews TD, Barnes C, Campbell P, Hu M, Ihm CH, et al. 2012. Origins and functional impact of copy number variation in the human genome. 464:704–712.

Cox DM, Butler MG. 2015. The 15q11.2 BP1-BP2 microdeletion syndrome: A review. Int J Mol Sci 16:4068–4082.

Fromer M, Purcell SM. 2014. Using XHMM Software to Detect Copy Number Variation in Whole-Exome Sequencing Data. Curr Protoc Hum Genet 81:7.23.1–7.23.21.

Gilissen C, Hehir-Kwa JY, Thung DT, Vorst M van de, Bon BWM van, Willemsen MH, Kwint M, Janssen IM, Hoischen A, Schenck A, Leach R, Klein R, et al. 2014. Genome sequencing identifies major causes of severe intellectual disability. Nature 511:344–347.

Glessner J, Bick AG, Ito K, Homsy J, Rodriguez-Murillo L, Fromer M, Mazaika EJ,Vardarajan B, Italia MJ, Leipzig J, DePalma S, Golhar R, et al. 2014. Increased frequency of de novo copy number variations in congenital heart disease by integrative analysis of SNP array and exome sequence data. Circ Res.

Homsy J, Zaidi S, Shen Y, Ware JS, Samocha KE, Karczewski KJ, Depalma SR, Mckean D, Wakimoto H, Gorham J, Jin SC, Deanfield J, et al. 2015. De novo mutations in congenital heart disease with neurodevelopmental and other congenital anomalies. Science (80-) 350:1262–1266.

Iossifov I, O’roak BJ, Sanders SJ, Ronemus M, Krumm N, Levy D, Stessman HA, Witherspoon K, Vives L, Patterson KE, Smith JD, Paeper B, et al. 2014. The contribution of de novo coding mutations to autism spectrum disorder. Nature 13:216–221.

Jeffares DC, Jolly C, Hoti M, Speed D, Shaw L, Rallis C, Sedlazeck FJ. 2017. Transient structural variations have strong effects on quantitative traits and reproduction isolation in fission yeast. Nat Commun 1–11.

Joshi RS, Garg P, Zaitlen N, Lappalainen T, Watson CT, Azam N, Ho D, Li X, Antonarakis SE, Brunner HG, Buiting K, Cheung SW, et al. 2016. DNA Methylation Profiling of Uniparental Disomy Subjects Provides a Map of Parental Epigenetic Bias in the Human Genome. Am J Hum Genet 99:555–566.

Jun G, Flickinger M, Hetrick KN, Romm JM, Doheny KF, Abecasis GR, Boehnke M, Kang HM. 2012. Detecting and estimating contamination of human DNA samples in sequencing and array-based genotype data. Am J Hum Genet 91:839–48.

Kloosterman WP, Francioli LC, Hormozdiari F, Marschall T, Hehir-kwa JY, Abdellaoui A, Lameijer E, Moed MH, Koval V, Renkens I, Roosmalen MJ Van, Arp P, et al. 2015. Characteristics of de novo structural changes in the human genome. Genome Res 792–801.

Layer RM, Chiang C, Quinlan AR, Hall IM. 2014. LUMPY: a probabilistic framework for structural variant discovery. Genome Biol 15:R84.

Mazaika E, Homsy J. 2014. Digital Droplet PCR: CNV Analysis and Other Applications. Curr Protoc Hum Genet 82:7.24.1–7.24.13.

McCarroll S a, Hadnott TN, Perry GH, Sabeti PC, Zody MC, Barrett JC, Dallaire S, Gabriel SB, Lee C, Daly MJ, Altshuler DM, The International HapMap Consortium. 2006. Common deletion polymorphisms in the human genome. Nat Genet 38:86–92.

Mccarroll SA, Hadnott TN, Perry GH, Sabeti PC, Zody MC, Barrett JC, Dallaire S, Gabriel SB, Lee C, Daly MJ, Altshuler DM, Hapmap I. 2006. Common deletion polymorphisms in the human genome. 38:86–92.

O’Roak BJ, Deriziotis P, Lee C, Vives L, Schwartz JJ, Girirajan S, Karakoc E, Mackenzie AP, Ng SB, Baker C, Rieder MJ, Nickerson D a, et al. 2011. Exome sequencing in sporadic autism spectrum disorders identifies severe de novo mutations. Nat Genet 43:585–9.

Quinlan AR, Clark RA, Sokolova S, Leibowitz ML, Zhang Y, Hurles ME, Mell JC, Hall IM. 2010. Genome-wide mapping and assembly of structural variant breakpoints in the mouse genome. Genome Res 623–635.

Sanders SJ, Murtha MT, Gupta AR, Murdoch JD, Raubeson MJ, Willsey AJ, ErcanS Sencicek AG, DiLullo NM, Parikshak NN, Stein JL, Walker MF, Ober GT, et al. 2012. De novo mutations revealed by whole-exome sequencing are strongly associated with autism. Nature 485:237–241.

Szatkiewicz JP, O’Dushlaine C, Chen G, Chambert K, Moran JL, Neale BM, Fromer M, Ruderfer D, Akterin S, Bergen SE, Kähler a, Magnusson PKE, et al. 2014. Copy number variation in schizophrenia in Sweden. Mol Psychiatry 19:762–73.

Tattini L, D’Aurizio R, Magi A. 2015. Detection of genomic structural variants from next generation sequencing data. Front Bioeng Biotechnol 3:.

Wala J, Bandopadhayay P, Greenwald N, Rourke RO, Stewart C, Schumacher S, Li Y, Weischenfeldt J, Nusbaum C, Campbell P, Meyerson M, Zhang Z. 2017. SvABA: Genome-wide detection of structural variants and indels by local assembly. bioRxiv 1–40.

Weischenfeldt J, Symmons O, Spitz F, Korbel JO. 2013. Phenotypic impact of genomic structural variation: insights from and for human disease. Nat Rev Genet 14:125–138.

Weisenfeld NI, Yin S, Sharpe T, Lau B, Hegarty R, Holmes L, Sogoloff B, Tabbaa D, Williams L, Russ C, Nusbaum C, Lander ES, et al. 2014. Comprehensive variation discovery in single human genomes. Nat Publ Gr 46:1350–1355.

Werling DM, Brand H, An J-Y, Stone MR, Glessner JT, Zhu L, Collings RL, Dong S, Layer RM, Markenscoff-Papadimitriou E, Farrell A, Schwartz GB, et al. 2017. Limited contribution of rare, noncoding variation to autism spectrum disorder from sequencing of 2,076 genomes in quartet families. bioRxiv 1–45.

Zook JM, Catoe D, Mcdaniel J, Vang L, Spies N, Sidow A, Weng Z, Liu Y, Mason CE, Alexander N, Henaff E, Mcintyre ABR, et al. 2016. Data Descriptor: Extensive sequencing of seven human genomes to characterize benchmark reference materials. Nature 1–26.

